# Transcriptome Analysis in Yeast Reveals the Externality of Position Effect

**DOI:** 10.1101/2020.04.02.021162

**Authors:** Qian Gui, Shuyun Deng, Wenjun Shi, Xiujuan Cai, Zhen-Zhen Zhou, Jian-Rong Yang, Xiaoshu Chen

**Author notes:** Correspondence should be addressed to X.C. or J.-R.Y. These authors contributed equally to this work (Qian Gui, Shuyun Deng).

## Abstract

When a gene is integrated into the chromosome, its activity depends on the genomic context. Although this phenomenon of “position effect” was widely reported, how the integration event affects the local environment, or the “externality” of position effect, remained largely unexplored, let alone the mechanism or phenotypic consequence of such externality. Here, we examined the transcriptome profiles of ∼250 *Saccharomyces cerevisiae* strains, each with *GFP* inserted into a different locus of the wild-type strain. We found that the GFP expression level and the change of expression of genes near the integration site decreases in genomic regions with high density of essential genes. This observation was found associated with H3K4me2 by further joint-analysis with public genome-wide histone modification profiles. More importantly, we found that the expression changes of neighboring genes, but not the GFP expression, exerted a significant impact on cellular growth rate. As a result, genomic loci that grant higher GFP expression immediately after the integration will have lower total yield of GFP in the long run. Our results, which were consistent with the competition for transcriptional resources among neighboring genes, revealed a previously unappreciated facet of the position effect, and highlighted its impact on the fate of genomic integration of exogenous genes, which has profound implications for biological engineering and pathology of virus integrative to host genome.

## Introduction

Gene integration into the genome is a major type of genomic alterations commonly observed in both natural (e.g., viral integration into host genome [1], transposon [2] and horizontal gene transfer [3]) and artificial circumstances [4]. Depending on the location of the genomic integration, the activity of the integrated gene varies substantially [5, 6], as does the phenotypic outcome of the integration event [7-9], a phenomenon commonly referred to as “position effect” [10]. A classic example of position effect is the translocation of white gene of fruit fly (*Drosophila melanogaster*) into heterochromatin, giving the original solid red eye a white and red mottled appearance [7, 8]. Recently, genome-wide studies have provided more mechanistic details on position effects, such as the regulatory role of enhancers, gene order, various epigenetic modifications, chromatin domains and three dimensional localization [5, 11-13]. It is therefore not surprising that position effects have a significant impact on the evolution of chromosome organization [14], improvements of genetic engineering [15] and a number of genetic diseases [9, 16].

However, despite extensive efforts to clarify the influence of position effects on the function of focally integrated gene, much less is known about how integration events affect other genes. A previous study has shown that integration into four different loci in yeast genome resulted in a few changes in the transcriptional profile [12]. In addition, the separate integration into the 63 loci of yeast chromosome 1 did not cause dramatic changes in the expression level of HO locus on chromosome 4 [17]. However, these results were derived from only on a few integration events or one distal gene, which might not be representative enough regarding the impacts for genes of different linear or three dimensional distance from the integration site.

Note that our focus was on general patterns independent of gene function, just as “position effect” is a general phenomenon regardless of the function of the integrated gene. Specifically, does the transcriptional activity of other genes in the genome change due to the integration event? If so, how are these changes influenced by the genomic location of the integration? Last but not least, do these changes contribute to the phenotypic consequence of position effect? Theoretically, integration of a gene would significantly alter the transcriptional regulation, thereby causing changes in activity of other genes, especially those share local transcriptional resources with the integrated gene. On the one hand, it is possible that the integrated gene competes with nearby genes for local transcriptional resources, thereby reducing the activities of nearby genes, such as in the phenomenon of promoter interference [18, 19]. On the other hand, it is also possible that the promoter of the integrated gene recruits more transcriptional resources, such that nearby genes gain access to more transcription resources, thereby improving their activity. These two possibilities are hereinafter referred to as “transcriptional competition” and “transcriptional synergism”.

In this context, it is commonly accepted that the order of genes in the genome and the transcription profiles of the wild-type strain are the result of long-term evolutionary optimization [20, 21]. In particular, one prevailing theory proposed that, essential genes, which cause lethality when deleted, tend to cluster on the genome, thereby forming open regions for continuous transcription. This non-random distribution of essential genes ensures that their expression noise, which is likely highly deleterious when happen to essential genes, is minimized [14, 17, 22]. Based on this theory, we speculated that the density of essential gene is a proxy for the allocation of intracellular transcriptional resources, and therefore dictates the prevalence of “transcriptional competition” or “transcriptional synergism”.

To test our hypothesis, we conducted transcriptome deep-sequencing for ∼250 *Saccharomyces cerevisiae* strains, which were randomly picked from a previously constructed library with GFP reporter cassettes individually inserted into various loci across all chromosomes [17]. We found that integration into genomic loci whose linear or three-dimensional proximity were enriched with essential genes decreased the transcriptional activity of the integrated gene as well as adjacent genes, which is consistent with the model of transcriptional competition. In contrast, integration into genomic loci where essential genes were depleted increased the transcriptional activity of the integrated and adjacent genes, which is consistent with the model of transcriptional synergism. The observed “externality” of position effect was at least partially explainable by H3K4me2 methylation of the surrounding genomic regions. Intriguingly, the expression changes of neighboring genes, rather than that of the integrated gene, was correlated with the rate of cellular growth, revealing a previously underappreciated mechanism for the phenotypic consequence of position effect.

## Results

### Transcriptome sequencing and quality control

We previously replaced the kanMX module in the heterozygous deletion strains of *S. cerevisiae* at hundreds of different loci across all chromosomes, with an expression cassette that comprises the marker gene URA3 and a GFP gene driven by the *RPL5* promoter (pRPL5-GFP) [17]. To examine the effect of this inserted GFP cassette on adjacent genes, we randomly selected over 250 strains from the reconstructed heterozygous deletion library and sequenced the transcriptome of each selected strain (see Methods, Supplemental Table S1 and S2). We further performed the following quality control steps to ensure the transcriptome datasets are comparable against each other. First, we confirmed that our experiments were highly reproducible with strong correlations between transcriptome profiles from different biological replicates (Fig. 1A). Second, strains with severe haploinsufficiency due to the heterozygous deletion of endogenous genes were removed, such that only strains with transcriptome profiles of limited deviation from the wild-type strain (Pearson’s Correlation Coefficient > 0.9) were kept for subsequent analysis (Fig.1B). Third, we estimated for the heterozygously deleted genes the ratio between its expression in the constructed *GFP* strains and that in the wild-type strain, which was expected as 0.5. A constructed strain was excluded from further analyses if this ratio appeared as an outlier among all the constructed strains (Fig.1C), which ensured negligible effect of feedback regulation over the heterozygously deleted gene. Finally, the transcriptome profiles of the constructed strains representing 240 loci inserted with the GFP cassette were retained for downstream analyses.

**Figure 1.**
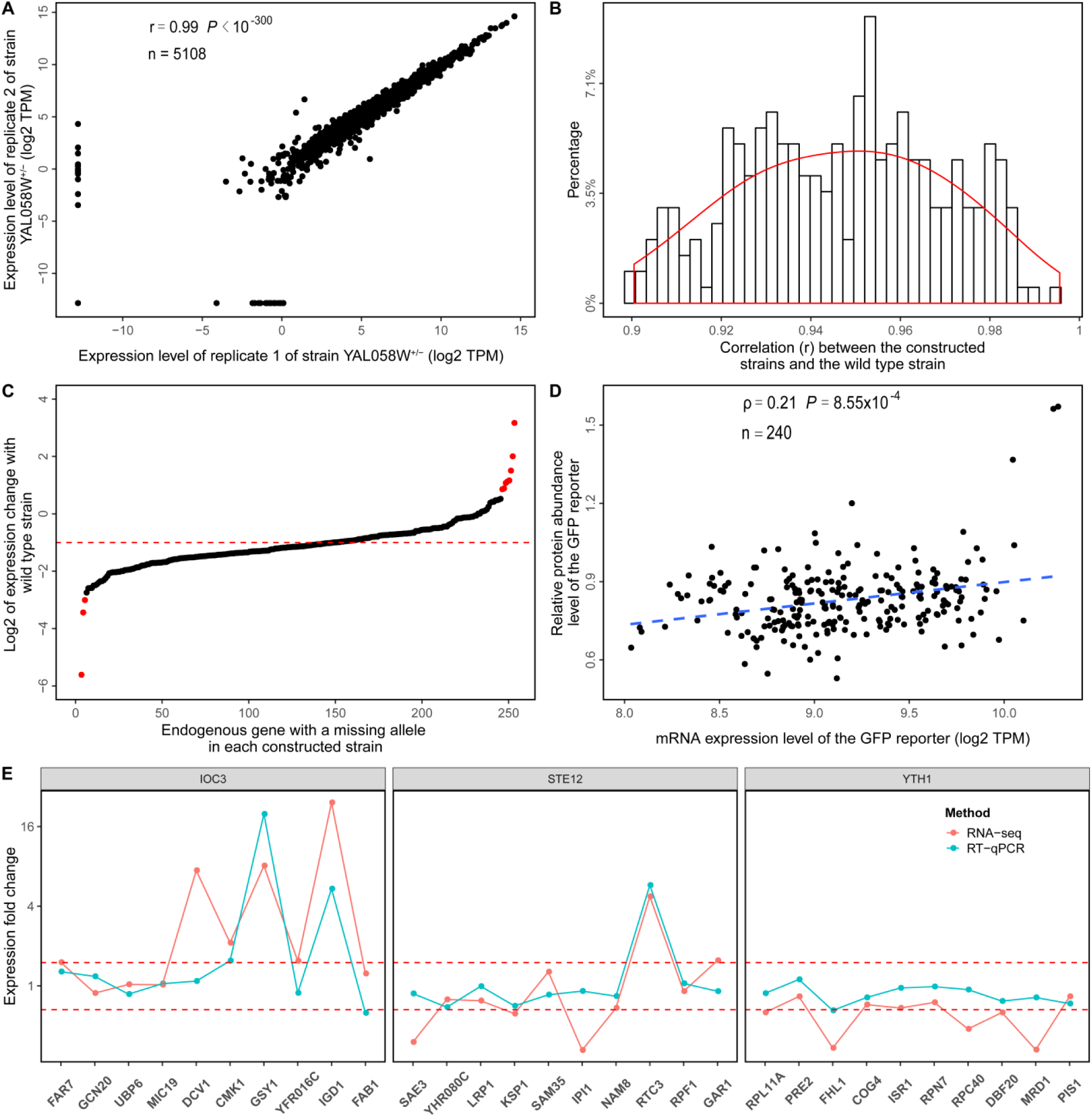
High quality of our transcriptome dataset. **(A)** Reproducibility between two biological replicas of strain YAL058W^+/−^. **(B)** Distribution of the Pearson’s correlation coefficient between the transcriptome profile of each constructed strain and the transcriptome profile of the wild-type strain. The red line represented the kernel density estimation of the distribution of the correlation coefficient. **(C)** Expression changes of endogenous gene missing one allele in each constructed strain compared to the wild-type strain. The red dots indicate the constructed strains whose expression changes of endogenous gene are outliers of the expression change of all examined endogenous genes missing one allele. The red dotted line indicates the expected value of 0.5. **(D)** The mRNA expression level of the GFP reporter we tested was significantly correlated with the protein abundance level measured in previous studies [17]. The dotted line represents fitted linear regression model. **(E)** The expression fold change compared with the wild-type strain in 5 genes each upstream and downstream (*x* axis) of the GFP insertion site of three constructed strains detected by RT-qPCR and RNA-seq.

To further corroborate the reliability of our transcriptome datasets, we compared the RNA-seq-based mRNA expression levels of GFP with previously measured protein abundance levels based on flow-cytometry [17]. We found that the mRNA expression is significantly correlated with the protein abundance (Spearman’s Rank Correlation Coefficient ρ = 0.21, *P* < 9 × 10^−4^, Fig.1D). Moreover, the mRNA expression spanned a five-fold range, whereas the protein abundance spanned a 2.5-fold range (Fig.1D). Such slight decrease in range of expression in protein abundance relative to that of mRNA expression is also consistent with known translational buffering of the transcriptional variation in yeast [23]. In addition, the previously observed effect of chromatin status [12], such as H3K4me1 modification, is also significant for the mRNA expression of GFP and the native gene (Fig.S1). Furthermore, we selected three constructed strains and quantified the mRNA expression levels of ten genes flanking the GFP insertion site (five on each side) by RT-qPCR. We found that the fold changes of expression relative to the wild-type strain as measured by RT-qPCR are highly consistent with that measured by RNA-seq (Fig.1E). Altogether, the above results indicated that our transcriptome dataset was of high quality, and thus a reliable dataset for the detection of the position effect and its potential externality.

### The externality of position effect within linear proximity

According to a widely accepted model explaining how position effect drives the nonrandom distribution of genes along the chromosomes, clusters of essential genes signify the genomic region where genes could have higher transcriptional activity and lower expression stochasticity [14, 17, 22]. In support of this model, we found in the transcriptome profile of the wild-type strain that the expression level of the surrounding genes was significantly correlated with the density of essential genes in the surrounding genomic region (Fig.2A and B). This observation is also consistent with the expression similarity commonly observed for neighboring genes [24].

However, when we examined the expression level of GFP in the heterozygous deletion strains, we were surprised to notice an opposite trend, where the GFP expression level was anti-correlated with the density of essential genes (Fig.2C). Furthermore, the GFP expression level is not related to the expression level of surrounding genes in the wild-type strain (ρ = -0.09, *P* = 0.15).

How to explain the opposite observations made with surrounding endogenous genes and the inserted GFP gene? The major difference between GFP and the endogenous genes were that the localization of the endogenous genes was presumably optimized by naturel selection, such that endogenous genes with promoters driving high and stable expression will likely be located in a genomic region enriched of essential genes. Whereas for the GFP, the *RPL5* promoter might face strong transcriptional competition when the local density of essential genes was high, since essential genes should have been evolved to be transcriptionally more competitive than non-essential genes because lowered expression of essential genes are generally more detrimental than non-essential genes.

To test whether genomic regions with higher density of essential genes indeed have stronger transcriptional competition, we examined the change of expression levels due to GFP integration for the genes surrounding the integration site. We found that as the local density of essential genes increases, the fraction of up-regulated genes decreases (Fig.2D), while the fraction of down-regulated genes increases (Fig.2E). The above observations remained qualitatively unchanged with different threshold for differential expression (Fig.S2). Moreover, the median fold change of the expression was anti-correlated with the local density of essential genes in the genomic region surrounding the integration site (Fig.2F). As a result, GFP integration at a genomic region with high local density of essential genes led to significant down-regulation of genes surrounding the integration site. On the contrary, the expression level of genes surrounding the integration site will increase only at genomic regions with no adjacent essential gene. In other words, transcriptional synergism caused by the inserted highly expressed gene was applicable only when there was no adjacent essential gene. Therefore, these results suggested a stronger transcriptional competition in genomic regions with higher local density of essential genes.

Notwithstanding, since the expression changes of the surrounding genes was apparently not caused by the function of GFP, the above observation therefore suggested that the genomic context of gene integration not only affects the transcriptional activity of focal integrated gene, but also that of the surrounding genes. Such externality constituted a previously unrecognized facet of the position effect. To determine the range of this externality, we examined different spans of the genomic region in terms of genes flanking each side of the integrations site. We found that in the genomic region of up to 40 surrounding genes (20 on each side of the integration), the local density of essential genes were still anti-correlated with the GFP expression level, with the strongest correlation found at 18 surrounding genes (Fig.2G). For the fraction of up-regulated genes (Fig.2H), fraction of down-regulated genes (Fig.2I), and the median expression change of the endogenous genes (Fig.2J), the span of genomic region with a significant externality were up to 30, 40 and 20 surrounding genes, with the most significant externality effect appeared at 10, 28 and 16 surrounding genes, respectively. We also tried similar analyses using genomic distance measured by *k*ilo *b*ase *p*air (kbp) rather than the number of adjacent genes, which gave rise to similar patterns (Fig.S3).

Collectively, these results suggested that the externality of position effect could potentially influence the expression levels of tens of genes surrounding the integration site, presumably via competition for transcriptional resources. This phenomenon therefore deserves further investigation for its underlying mechanism and its contribution to the phenotypic consequence of position effect.

### The externality of position effect within three-dimensional proximity

The transcriptional resources were allocated not only linearly along DNA molecule, but also three-dimensionally in so-called “transcriptional factories” as different genomic regions fold into specific focal sites of active transcription [25]. If the externality of position effect can indeed be explained by the competition of transcriptional resource, we should predict similar effect for the density of essential genes in the three-dimensional proximity of the GFP integration site (Fig.3A). We thus tested the above prediction on the basis of a three-dimensional model of the yeast genome [26].

We found patterns similar to those observed in the linear proximity of the GFP integration site, which were listed as follows. First, in a three-dimensional proximity enriched with essential genes, the expression of the adjacent genes in the wild-type strain tended to be higher (Fig.3B), but the integrated GFP in modified strains tended to be lower (Fig.3C). Second, the density of essential genes in the three-dimensional proximity was negatively correlated with the fraction of up-regulated genes (Fig.3D), positively correlated with the fraction of down-regulated genes (Fig.3E), and negatively correlated with the median expression change of genes within the three-dimensional proximity. Third, the observed externality of position effect remained significant if we considered different number of genes (Fig.3G-J), or physical distance (in *n*ano*m*eter, or nm, Fig.S4) within the three-dimensional proximity of the GFP integration site. Fourth, although majority (∼80%) of the three-dimensional adjacent genes were also linear neighbors of the GFP integration site, excluding these linear neighbors did not change our conclusion on the significance of externality of position effect (Fig.S5). Altogether, these results again supported the externality of position effect, and suggested that such externality could be explained by the competition for transcriptional resources impacted by the integration event, therefore expanding our understanding of position effects.

### Contribution of histone modifications to the externality of position effects

Since histone modifications can predict the expression of reporter genes [12], we hypothesized that histone modifications are also involved in regulating the expression changes around the integration site. We collected high-throughput sequencing-based profiles for eight major types of histone modifications (H3K4me1, H3K4me2, H3K4me3, H3K36me3, H3K79me3, H3K9ac, H3K12ac and H3K14ac. See supplemental Table S1 and S3) in the wild-type strain, estimated the histone modification levels for the genes around the integration site, and compared them with the local density of essential genes and the expression level of the integrated GFP (See Materials and Methods).

We found that in genomic regions with high expression level of GFP gene, the surrounding genes tend to have significantly high H3K4me1 and H3K4me2 signals (Fig.4A). Such patterns is consistent with the report that H3K4me1 and H3K4me2 were generally found as associated with active transcription [27]. In addition, we found that H3K4me2 signal was positively correlated with the fraction of up-regulated adjacent genes, negatively correlated with the fraction of down-regulated adjacent genes, and positively correlated with the median fold change of expression of the adjacent genes (Fig.4B). Analyses for three-dimensional proximity of the integration site also confirmed the above conclusion (Fig.S6). These results suggested that H3K4me2 signal could be used as a marker of the expression level of integrated gene and the adjacent genes.

However, H3K4me1 and H3K4me2 signals are significantly negatively correlated with the density of essential genes (Fig.4A), which are considered to be at high expression levels. These observations indicated functional inter-relationships among the aggregation of essential genes, the presence or absence of particular histone modifications and the high transcriptional competition in the region. We then used partial correlation analysis to find out which of the three factors is decisive. As for the GFP expression level, we found that it was not correlated with H3K4me2 when essential gene density is controlled (Fig.4C), and it was anti-correlated with essential gene density when H3K4me2 level was controlled (Fig.4D). These patterns suggested that given the transcriptional activity of the focal site, the essential gene density in the neighborhood determined the final expression level of the integrated gene, which was compatible with the model of transcriptional competition underlying position effect.

As for the externality of the position effect, we found that H3K4me2 level was still positively correlated with median expression changes of the genes within the surrounding genomic region after controlling the local density of essential genes (Fig.4C). Conversely, controlling H3K4me2 level will yield a non-significant correlation between density of essential genes and median expression change of adjacent genes (Fig.4D). These results suggested that the externality of position effect can at least be partially explained by the native level of H2K3me2 modification level, which was presumably evolutionarily optimized to fit the local density of essential genes, thereby creating a correlation between density of essential genes and expression of neighboring genes.

### The Position effect of the reporter gene on the fitness of the strain

With the externality of position effect, it is necessary to investigate the contribution of such externality to the phenotypic consequence of position effect. We chose to measure the single most important phenotype of yeast, i.e. fitness, by measuring the growth curve of each constructed strain in the rich medium YPD (see Methods). Surprisingly, unlike previous reports using multiple copies of integrated gene [28, 29], we found negligible effect of the GFP expression on the fitness (Fig.5A), possibly explainable by the relatively small variation for the single copy GFPs integrated in different loci. Meanwhile, we observed a signifi negative correlation between changes in the expression of adjacent genes and fitness (Fig.5B), indicating that the externality of position effect can have a major role in the phenotypic consequence of position effect.

How will the externality of position effect impact the evolutionary fate of integrated gene? On the one hand, the integrated gene (GFP in our system) would be transcribed to a higher abundance if it was inserted to a genomic region depleted of essential genes (Fig. 2C and Fig. 3C). On the other hand, the externality of such position effect dictated that the adjacent genes would be more likely up-regulated (Fig.2F and Fig.3F), thereby decreasing the cellular fitness to a greater extend (Fig.5B). As a result, the total transcriptional yield of GFP by the whole population might be higher for strains expressing GFP in regions depleted of essential genes (Fig.5C). But in the long run, when the growth rate of the population (fitness) is taken into consideration, the strains expressing GFP in regions enriched of essential genes might eventually transcribe GFP mRNA to a higher total amount (Fig.5C). Analyses for three-dimensional proximity of the integration site also support the above conclusion (Fig.S7). Theoretically, this feature should give rise to a trade-off between the immediate expression of the inserted foreign genes and its long-term population-wide transcriptional yield via lowered fitness of the cells with the inserted gene, which was mediated by the externality of the position effect (Fig.5B), but not by the expression of the focal inserted genes (Fig.5A).

**Figure 2.**
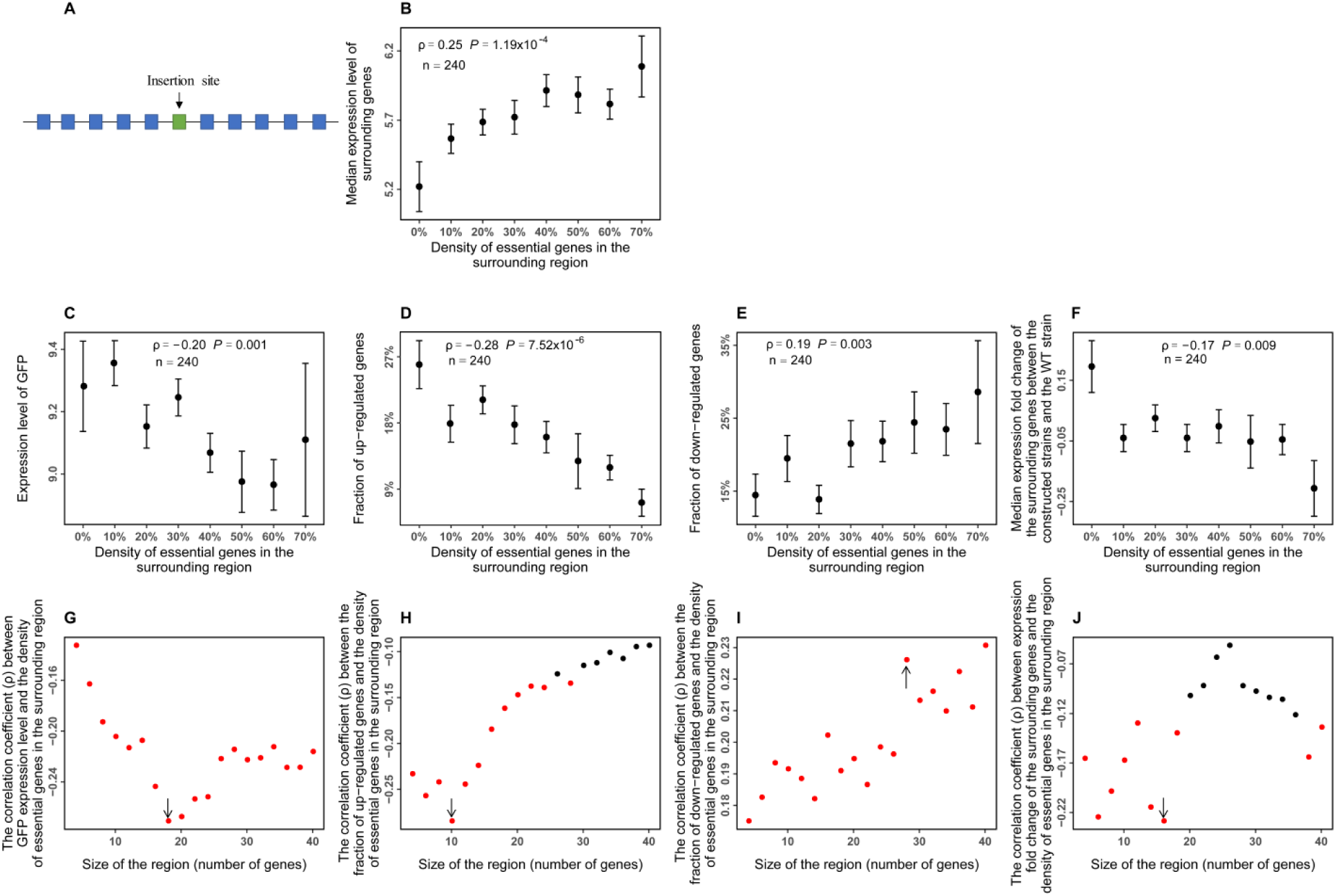
The expression level of the GFP gene and the changes in the expression level of adjacent genes are related to the content of the essential gene near the integration site. **(A)** Linear positional relationship between GFP integration site and adjacent genes. **(B-F)** The median expression level of adjacent genes before integration (B), the expression level of GFP reporter after integration (C), the fractions of up-regulated genes (D), the fractions of down-regulated genes (E), and the median expression fold changes of adjacent genes (F) were significantly related to the density of essential genes in the surrounding region. The surrounding regions included 5 genes each upstream and downstream of the GFP insertion site. The points represent the mean and the error bars represent the standard error within each range of essential gene density. **(G-J)** The correlation coefficient with the density of essential genes were shown for the expression level of GFP (G), the fractions of up-regulated genes (H), the fractions of down-regulated genes (I) and the median expression fold changes of adjacent genes (J) when genomic regions of different sizes (in terms of number of adjacent genes) were considered. The sizes of the region represent the total number of upstream and downstream adjacent genes. Compared with the wild strain, genes in the constructed strains whose expression level is increased to more than 1.5 fold were considered to be up-regulated, and genes whose expression level is reduced to less than 66.7% were considered to be down-regulated. The red dots indicate that the *P* values of the correlation coefficient are smaller than 0.05 (i.e., statistically significant), and the black dots indicate that they are greater than or equal to 0.05 (i.e., statistically not significant).

**Figure 3.**
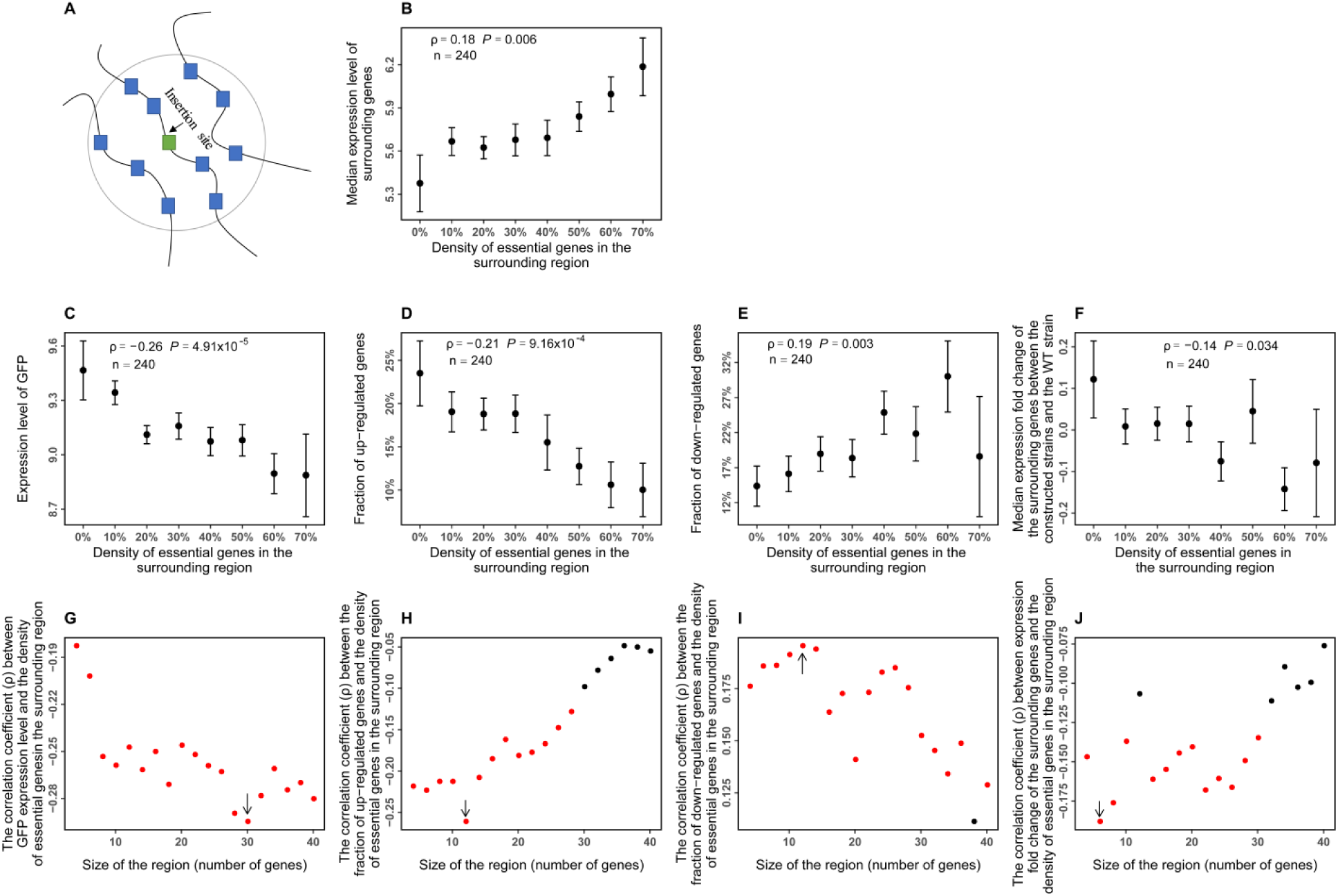
The expression level of the GFP gene and the changes in the expression level of neighboring genes are related to the content of the essential gene in the three-dimensional proximity of the integration site. **(A)** Three-dimensional proximity of the GFP integration sites and neighboring genes. **(B-F)** The median expression level of neighboring genes before integration (B), the expression level of GFP reporter after integration (C), the fractions of up-regulated genes (D), the fractions of down-regulated genes (E), and the median expression fold changes of neighboring genes (F) were significantly related to the density of essential genes in the three-dimensional proximity (including ten genes) around the integration site. The points represent the mean and the error bars represent the standard error within each range of essential gene density. **(G-J)** The correlation coefficient with the density of essential genes were shown for the expression level of GFP (G), the fractions of up-regulated genes (H), the fractions of down-regulated genes (I) and the median expression fold changes of neighboring genes (J) when genomic regions of different sizes (in terms of number of adjacent genes) were considered. The sizes of the region represent the total number of genes in three-dimensional proximity to the GFP integration site. The red dots indicate that the *P* values of the correlation coefficient are smaller than 0.05 (i.e., statistically significant), and the black dots indicate that they are greater than or equal to 0.05 (i.e., statistically not significant).

**Figure 4.**
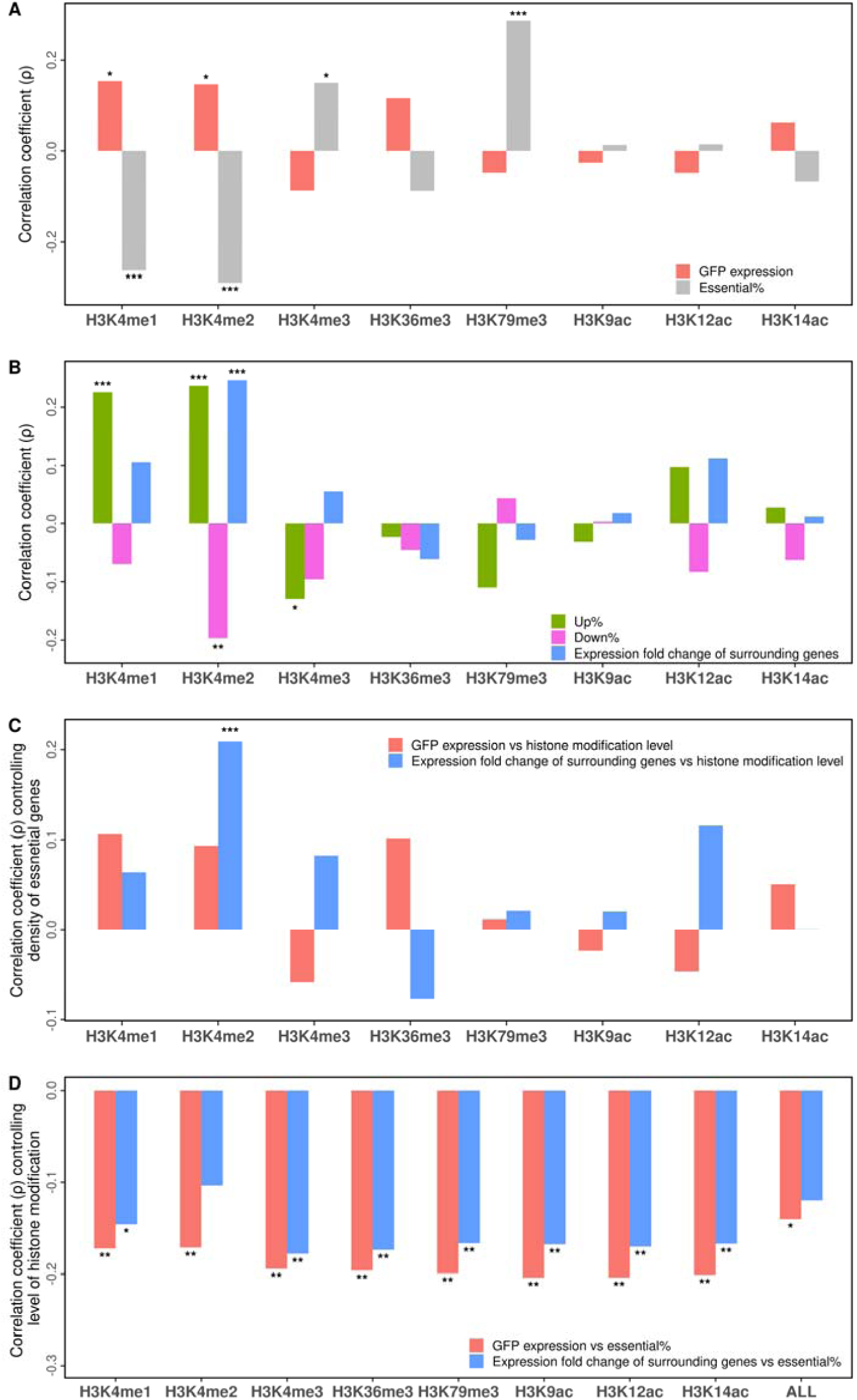
The expression level of the GFP gene and the changes in the expression level of adjacent genes are related to the H3K4me2 modification level of the surrounding region. **(A-B)** Correlations with the strengths of eight histone modifications of genes surrounding the integration site were shown for the expression level of GFP gene (A, red bars), the local density of essential genes (A, gray bars), the fractions of up-regulated genes (B, green bars), the fractions of down-regulated genes (B, pink bars), and the median expression fold changes of adjacent genes (B blue bars). **(C-D)** Partial correlations between the density of eight histone modifications of genes surrounding the integration site after controlling the local density of essential genes were shown for the expression level of GFP (C, red bars) and the median expression fold changes of adjacent genes (C, blue bars). Partial correlations with the local density of essential genes after controlling the levels of histone modifications were shown for with the expression level of GFP gene (D, red bars) and the median expression fold changes of adjacent genes (D, blue bars). The surrounding regions included 5 genes each upstream and downstream of the GFP insertion site. Statistical significance of the correlation was indicated: *: *P* < 0.05, **: *P* < 0.01, ***: *P* < 0.001.

**Figure 5.**
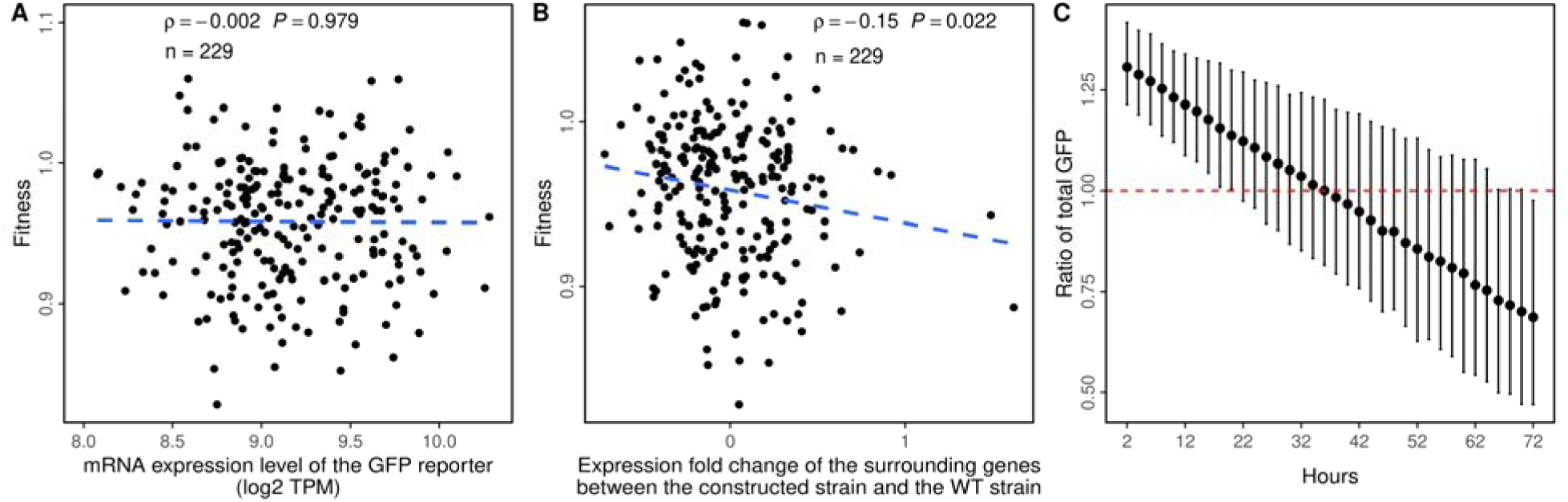
The changes in the expression level of adjacent genes, but not the expression level of the GFP gene, is related to the fitness of the constructed strains. **(A)** Correlation between the expression level of the GFP gene and the fitness of the constructed strains relative to the wild-type strain. **(B)** Correlation between the expression fold change of genes surrounding the integration site and the fitness of the constructed strains relative to the wild-type strain. **(C)** The ratio of total yield of GFP mRNA in the strains with GFP inserted to genomic regions with 10% adjacent essential genes, relative to that with 60% adjacent essential genes in different time-point of cultivation (from 2 hours to 72 hours, assuming a generation time of 1.5 hour). The 95% confidence interval of this ratio was estimated by bootstrapping the genes 1,000 times.

## Discussion

In this study, we determined the transcriptome profiles of ∼ 250 yeast strains, each with a GFP expression cassette integrated to a different locus of the genome. We found that the GFP expression levels were negatively correlated with local density of essential genes within either linear or three-dimensional proximity of the integration site. An opposite trend for the expression level of the neighboring genes around the integration site was also revealed, indicating previously unappreciated externality of position effect. Assuming that essential genes were transcriptionally more competitive, the observed position effect and its externality can be explained by competition between adjacent genes for transcriptional resources. We also found that specific histone modifications were closely related with the position effect and its externality. More importantly, the observed externality, but not the expression of the inserted gene, seemed to be responsible for the phenotypic consequence of the position effect. Altogether, our results revealed a previously unappreciated facet of the position effect, which might have profound implications synthetic biology and evolutionary biology, such as genetic engineering that aimed at maximizing transcriptional yield of exogenous genes, as well as understanding the evolutionary forces behind gene orders/distributions on chromosomes.

There were a few potential caveats in our study that worth discussion. First, yeast contains more than 4,000 verified genes, but only ∼250 gene loci have been tested in our analyses. Although our sampled loci were likely unbiased, future larger scale studies covering more loci should be carried out to examine the generality of our conclusion. Second, we used the promoter of *RPL5* (ribosomal 60S subunit protein L5), which is an essential gene, to drive the expression of GFP, and it was possible that the phenomenon presented here was promoter-specific. Third, the three-dimensional model of yeast chromosomes we used was measured in haploid strain rather than in the diploid strain, although dramatic differences in three-dimensional genomes of haploid and diploid cells were unlikely [30]. In addition, if the three-dimensional genomes of haploid and diploid cells were totally different, we should not be able to identify any statistical significance using the three-dimensional proximity based on the haploid cells.

Results from our study highlights how the evolutionary fate of an exogenous gene integrated into the genome will be affected by the density essential genes near the integration site. On the one hand, if the integration event happened at a locus of high density of essential genes, the expression of the integrated gene and the neighboring genes might be lowered due to the strongly competitive transcriptional environment created by adjacent essential genes, meanwhile the cellular fitness was not much influenced. On the other hand, if the integration happened at a locus of low density of essential genes, the integrated gene and its neighbors might have higher expression, but the cellular fitness will be significantly decreased. Most importantly, the expression changes of neighboring genes have a non-negligible effect on the fitness impact of position effect, a novel phenomenon here termed as the externality of position effect.

One common type of exogenous gene integration that occurs naturally is the integration of viral genes into the host genome. Previous studies mostly only focus on the direct functional consequence of such integration [1], such as intergenic integration destroying cis-regulatory elements, or intra-genic integration intervening transcription or even the structure of endogenous genes. Our study suggested that integration of a transcriptionally active gene can at least impact tens of genes near the integration site, which will likely further influence host cellular fitness.

Such potential indirect functional consequence of the viral integration warrants further investigation.

## Methods

### RNA extraction and sequencing

Each strain of the pRPL5-GFP cassette was inoculated into 5 ml of YPD medium (1% yeast extract, 2% peptone, 2% glucose), then cultured overnight at 30 °C and 250 rpm. The saturated culture was then returned to OD660 = 0.2 in 4 mL YPD and growth continued at 30 °C until OD660 = 0.65-0.75. Total RNA was then extracted from cell lysates using the RNeasy Mini Kit (Qiagen) according to the manufacturer’s instructions. The quality of RNA was determined by A260/A230 ratio, A260/A280 ratio and concentration using NanoDrop. An A260/A230 ratio > 2 and an A260/A280 ratio of the range of 1.8 to 2.2 were considered acceptable. Finally, they were sequenced in paired-end mode using HiSeq 4000 (Illumina) with a read length of 150 bp (Supplemental Table S1). In addition, we used a PrimeScript RT reagent Kit with gDNA Eraser (TAKARA) to reverse transcribe 1ug of total RNA into cDNA according to the manufacturer’s instructions

### Calculation of RNA abundance

Yeast (*Saccharomyces cerevisiae*) S288C reference genome version R64-2-1 and corresponding genome annotation were obtained from the Saccharomyces Genome Database (SGD) [31]. To estimate the RNA abundance in each strain, we mapped the adaptor-trimmed and quality-filtered [32] short reads to the reference genome by HISAT2 [33]. Then *t*ranscripts *p*er *m*illion reads (TPM) [34] was estimated by StringTie [35] based on the mapping results, and was then used as the gene expression levels (Supplemental Table S2).

### RT-qPCR primer design and measurement

We searched the cDNA sequences of 30 related genes and the control gene Actin (ACT1) from the S288C reference genome version R64-2-1, and used the NCBI Primer BLAST to design RT-qPCR primers. The RT-qPCR products were limited to 100-200 bp, melting temperature of each RT-qPCR primer was between 58°C to 62°C (Supplemental Table S4). The concentration of cDNA (0.2 μL) were then measured by RT-qPCR with 10μM of forward and reverse primers each in 96-well plates, using iTaq Universal SYBR Green supermix (BIO-RAD) on a Roche LightCycler 96 Real-Time PCR Cycler machine. The cycling parameters for amplification are: 95°C for 30 seconds and 40 cycles of 95°C for 5 seconds and 60°C for 30 seconds. Signals were normalized to that of Actin and quantified by the ΔΔCt method [36]. The resulting expression levels were represented as mean ± S.D. of four independent experiments each ran in triplicates.

### Determination of linear gene cluster

According to yeast S288C genome annotation, 5108 verified genes were selected for subsequent data analysis. First, we grouped the yeast gene components into overlapping windows of a specific number of consecutive genes, with a step size of 1 and the window size of 2, 4, 6, … or 40. Subsequently, we counted the fraction (density) of essential genes in each window, the median expression level of genes in this window for wild-type strains, and the fraction of genes that were up-regulated (greater, or > 1.5 fold, or 2 fold) and down-regulated (less, or < 66.7%, or 50%), as well as the median change in gene expression levels within that window after inserting the GFP gene. Similarly, we set the yeast genome as overlapping windows of a specific number of base pairs (the step size was equal to 5 kbp and the window sizes were set to 10 kbp, 15 kbp, 20 kbp, … or 80 kbp) to calculate the above properties again.

### Determination of three-dimensional gene cluster

We used the haploid yeast 3D chromosomal architecture that was inferred through a chromosome conformation capture-on-chip (4C) coupled with massively parallel sequencing [26]. A file containing a list of chromosomal interactions identified from HindIII libraries was downloaded from the original report to infer the spatial distance between gene pairs. Each gene window was defined as a fixed number of genes (set to 2, 4, 6, … or 40, respectively) that are closest (with highest interaction probability) to the focal gene. Similar to the analysis at the linear level, we calculated the density of essential genes, the median expression level of genes in wild-type strains, the fraction of genes that were up-regulated and down-regulated, and the median change in gene expression levels within each window after inserting the GFP reporter gene. Besides, we set the distance from the GFP insertion site to 1, 2, 3, … or 15 nanometer respectively as the window size to calculate the above properties again.

### Calculation of histone modifications

We downloaded high-throughput sequencing data in wild-type yeast on 8 types of histone acetylation and histone methylation (H3K4me1, H3K4me2, H3K4me3, H3K36me3, H3K79me3, H3K9ac, H3K12ac and H3K14ac) on Sequence Read Archive (SRA) (Supplemental Table S1). By applying a pipeline similar to the one we used to quantify RNA abundance, we calculated the histone modification levels (i.e., abundance of short reads from high-throughput sequencing targeting specific modifications) of the 200 bp region starting 200 bp upstream of the start codon of each endogenous gene (Supplemental Table S3). We then calculated the average level of histone modifications in the 11 gene window size of the linear level and the three-dimensional level.

### Fitness measurement

To measure the growth rate of the GFP constructed strains, we cultured the cells in YPD medium at 30 °C overnight, and then 5ul of the saturated culture was transferred to 145ul of YPD in 96-well plates. Each 96-well plate was shaken on a BioTek Epoch2 microplate reader at 30 °C for 12 hours with the OD600 signal reading every 10 minutes. After repeating the experiment for at least 3 times for each strain, we calculated the doubling time for each constructed strain (*t*_cs_) according to Murakami et al. [37]. Based on the comparison with the doubling time of the wild-type strain (*t*_wt_), the relative fitness (*w*) of each GFP constructed strain was calculated as:

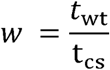

Additionally, the fitness was used to estimate the relative total yield of GFP mRNA between strains where GFP was integrated into loci with 10% essential genes compared to those with 60% essential genes by

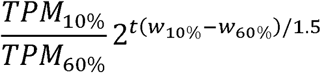

Where *TPM* was the mRNA expression level of GFP inferred from RNA-seq, *t* is time (in hours), and *w* is fitness.

## Acknowledgements

We thank Feng Chen and Xiaoyu Zhang for helpful discussions; and the anonymous reviewers for their insightful and helpful comments on this work. The overall research project was supported by grants from the National Special Research Program of China for Important Infectious Diseases (project number 2018ZX10302103, awarded to X. C), the National Key R&D Program of China (project number 2017YFA0103504, awarded to X. C, and project number 2018ZX10301402, awarded to J.-R. Y), the National Natural Science Foundation of China (project numbers 31771406 awarded to X. C, and 31671320, 31871320 and 81830103 awarded to J.-R. Y).

## Figures and legends

**Figure S1.**
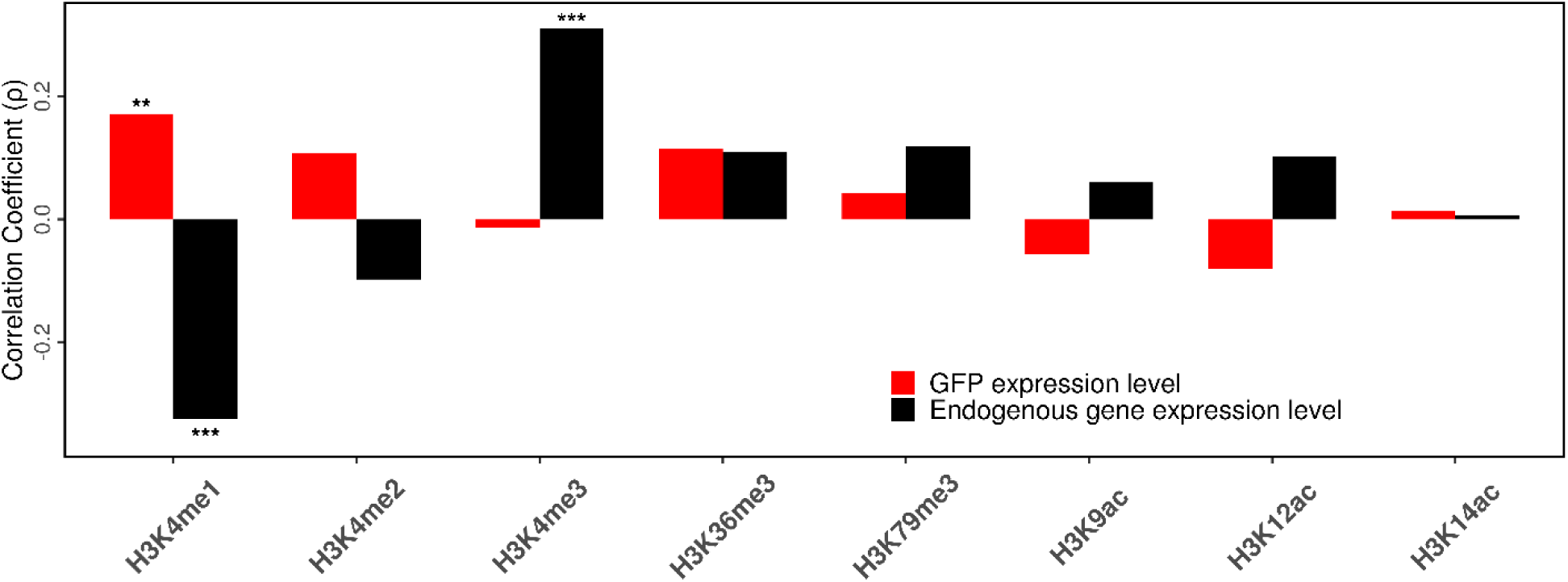
Correlations between the density of eight histone modifications in the integration site with the expression level of GFP gene (red bars) and the endogenous gene (black bars). Statistical significance of the correlation was indicated: *: *P* < 0.05, **: *P* < 0.01, ***: *P* < 0.001.

**Figure S2.**
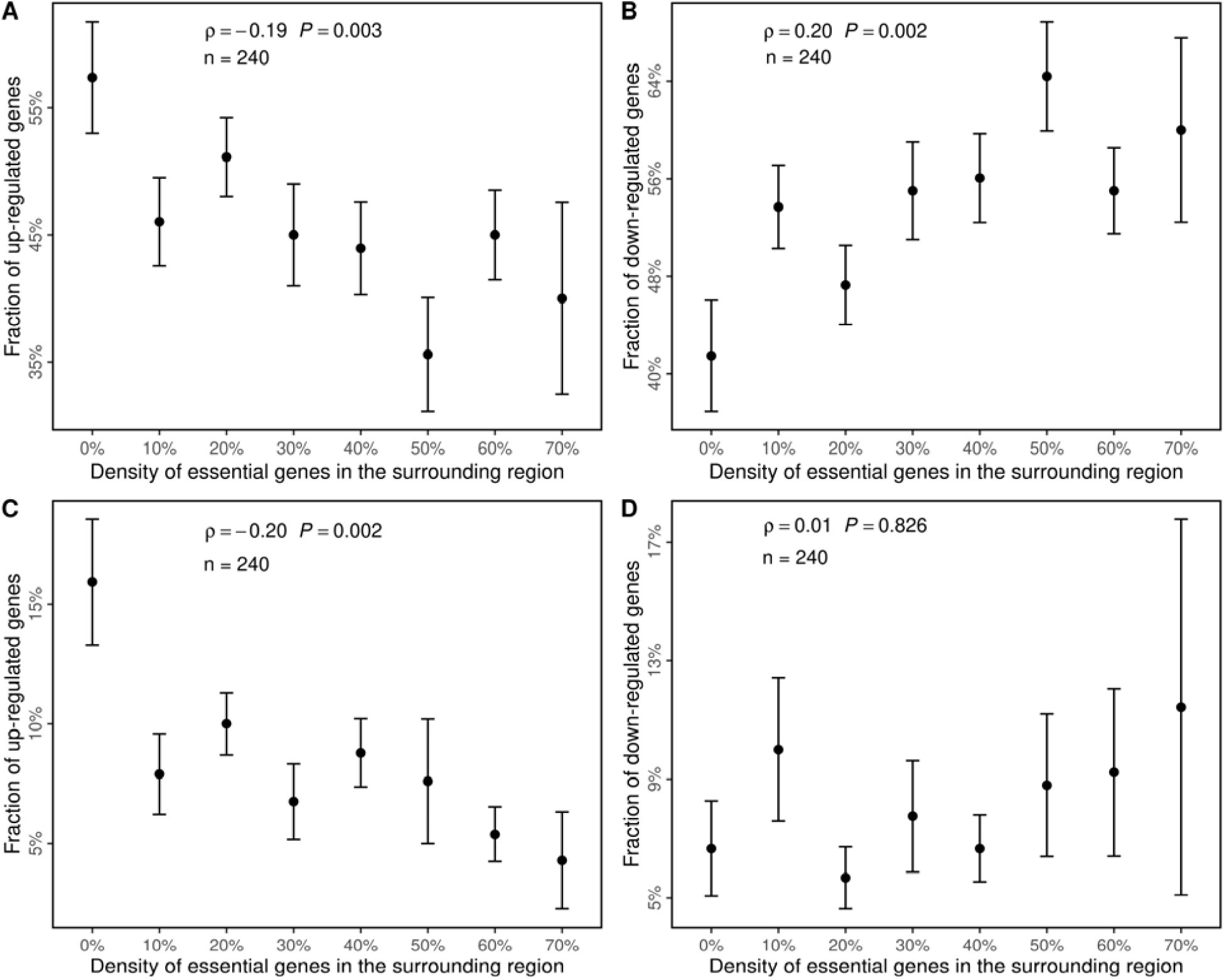
The changes in the expression level of adjacent genes are related to the content of the essential gene near the integration site. Similar to Fig 2D and E except that genes in the constructed strains whose expression level is greater than the wild-type strain were considered to be up-regulated (A), genes whose expression level is less than the wild-type strain were considered to be down-regulated (B), genes whose expression level is up-regulated by > 2 fold were considered to be up-regulated (C), and genes whose expression level is down-regulated to < 50% of the wild-type were considered to be down-regulated (D).

**Figure S3.**
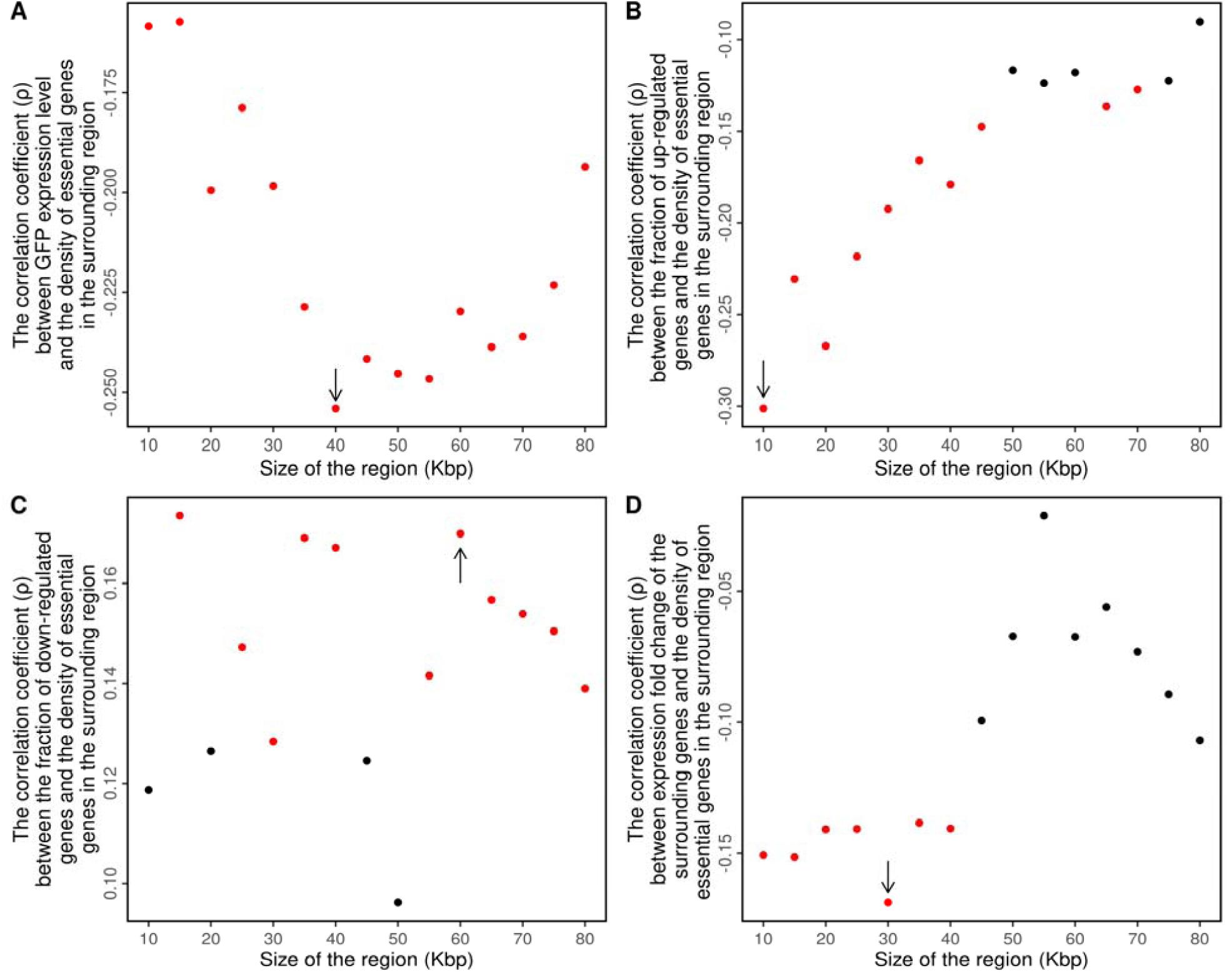
The expression level of the GFP gene and the expression fold changes of adjacent genes are related to the content of the essential gene near the integration site. Similar to Fig 2G to J except that the size of the region was measured by the distance (*k*ilo *b*ase *p*air, kbp) to the GFP integration site.

**Figure S4.**
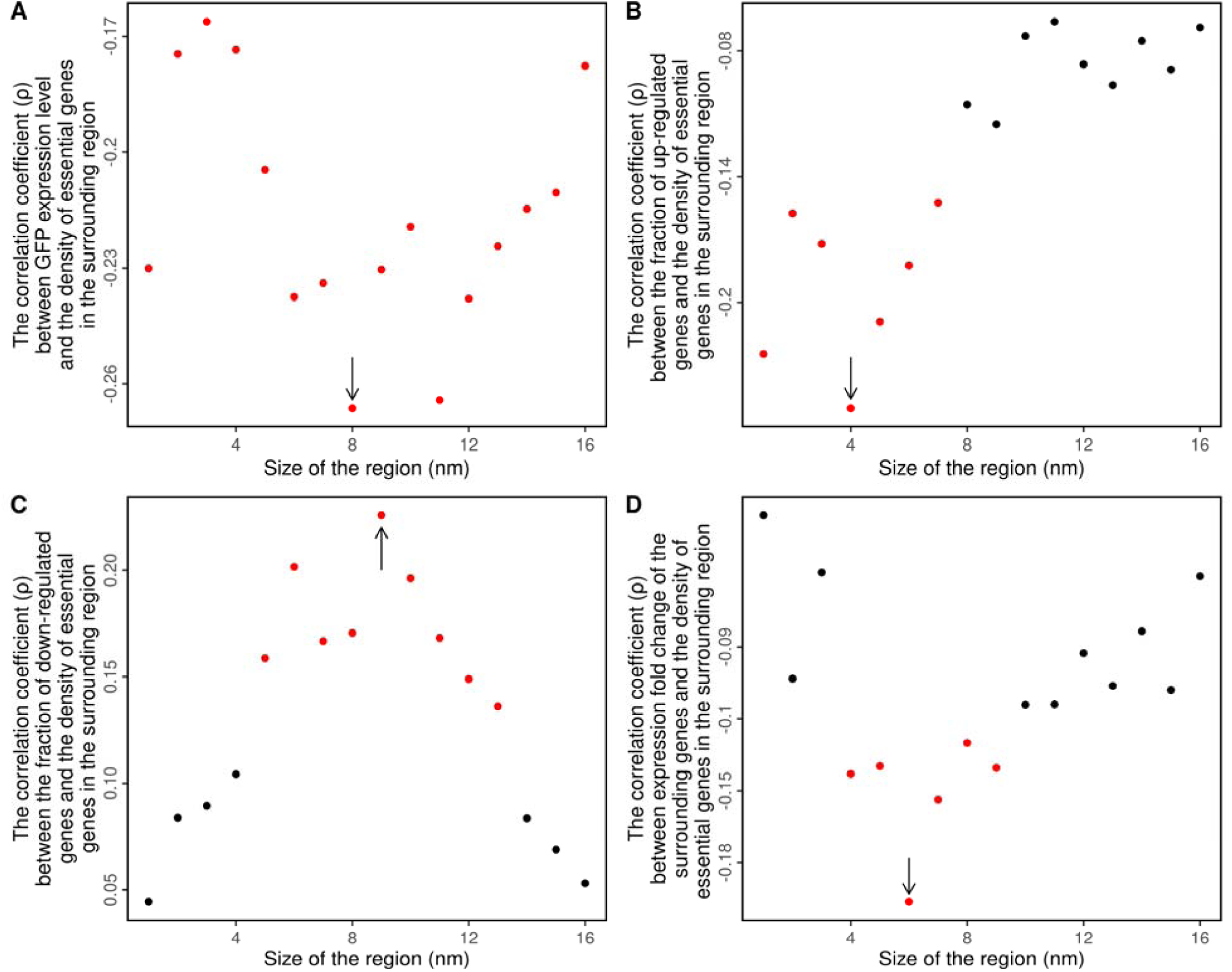
The expression level of the GFP gene and the expression fold changes of neighboring genes are related to the content of the essential gene in the three-dimensional proximity of the integration site. Similar to Fig 3G to J except that the size of the region was measured by the distance (*n*ano*m*eter, nm) to the GFP integration site. 1nm in space is approximately equal to 6.4kbp.

**Figure S5.**
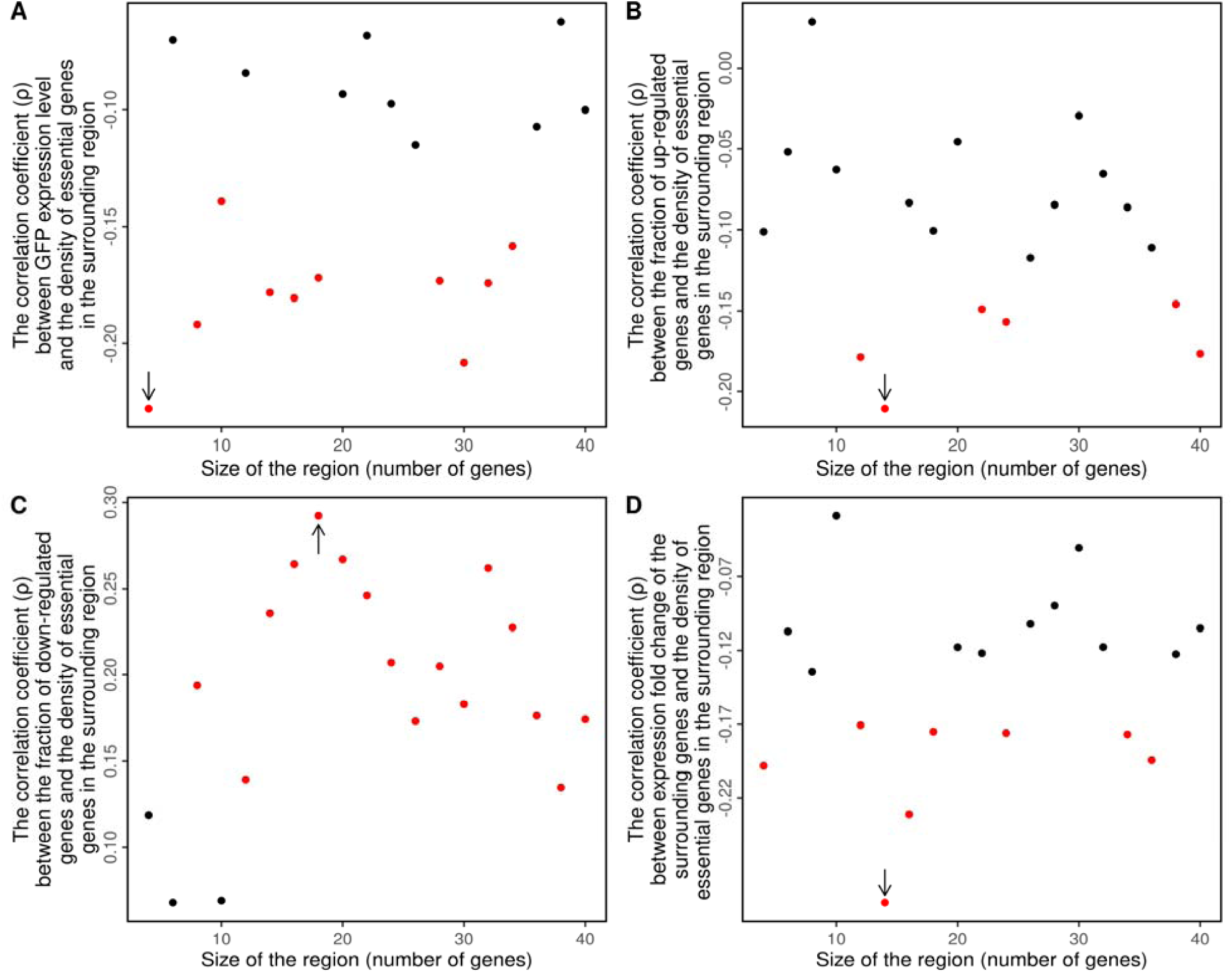
The expression level of the GFP gene and the expression fold change of neighboring genes are related to the content of the essential gene of the surrounding region in three-dimensional proximity but not in linear range. Similar to Fig 3G to J except that the surrounding area includes 10 genes that are spatially closest to the GFP insertion site but are not in the linear range.

**Figure S6.**
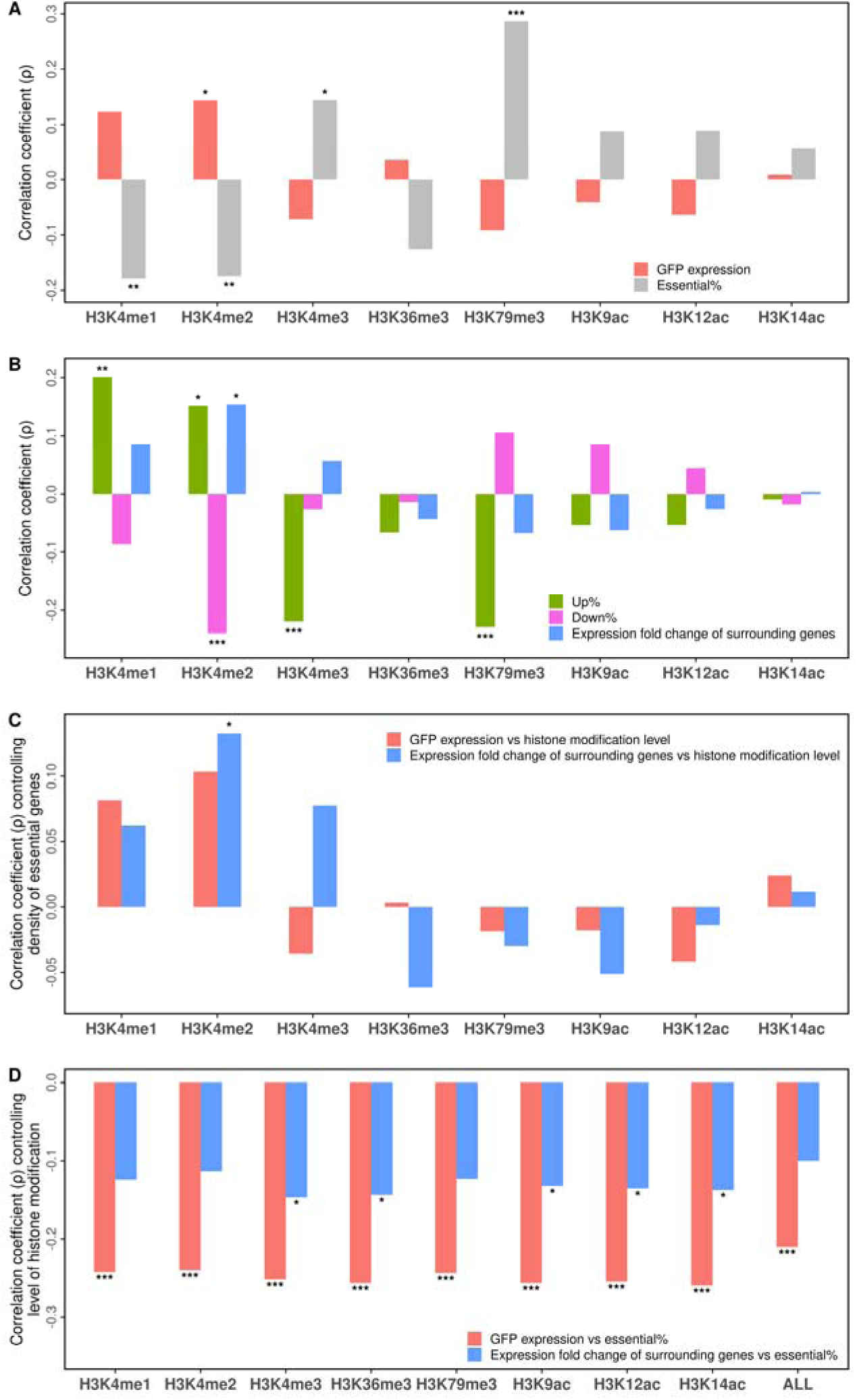
The expression level of the GFP gene and the expression fold change of neighboring genes are related to the level of H3K4me2 modification of the surrounding region in three-dimensional proximity. Similar to Fig 4 except that the surrounding region included the ten genes in closest proximity to the GFP insertion site.

**Figure S7.**
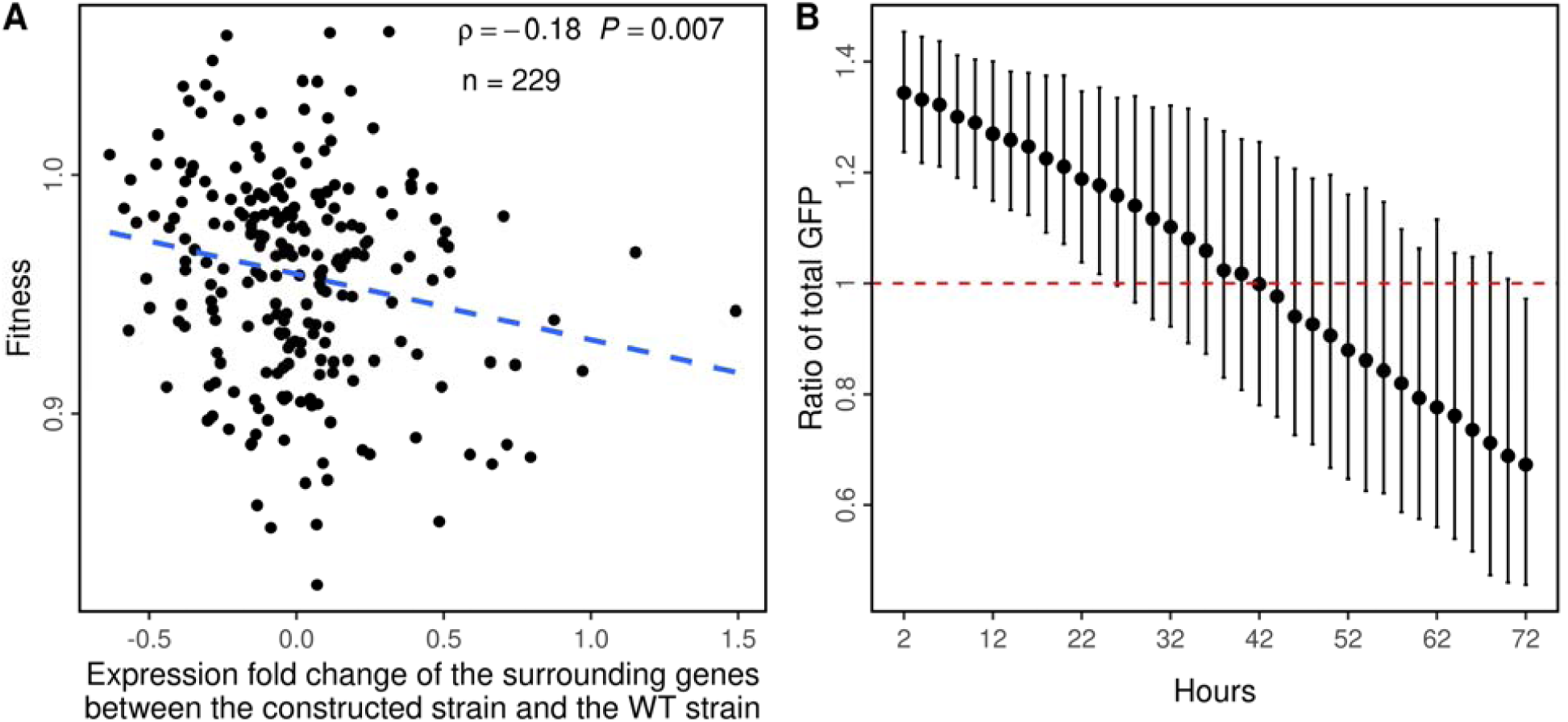
The changes in the expression level of neighboring genes, but not the expression level of the GFP gene, is related to the fitness of the constructed strains. Similar to Fig 5B to C, except that the surrounding region included the ten genes in closest three-dimensional proximity to the GFP insertion site.

## References

1. Ciuffi A: The benefits of integration. Clin Microbiol Infect 2016, 22:324–332.

2. Yant SR, Wu X, Huang Y, Garrison B, Burgess SM, Kay MA: High-resolution genome-wide mapping of transposon integration in mammals. Mol Cell Biol 2005, 25:2085–2094.

3. !!! INVALID CITATION !!!

4. Ivics Z, Li MA, Mates L, Boeke JD, Nagy A, Bradley A, Izsvak Z: Transposon-mediated genome manipulation in vertebrates. Nat Methods 2009, 6:415–422.

5. Akhtar W, de Jong J, Pindyurin AV, Pagie L, Meuleman W, de Ridder J, Berns A, Wessels LF, van Lohuizen M, van Steensel B: Chromatin position effects assayed by thousands of reporters integrated in parallel. Cell 2013, 154:914–927.

6. Feuerborn A, Cook PR: Why the activity of a gene depends on its neighbors. Trends Genet 2015, 31:483–490.

7. Sturtevant AH: The effects of unequal crossing over at the bar locus in Drosophila. Genetics 1925, 10:117.

8. Grewal SI, Jia S: Heterochromatin revisited. Nat Rev Genet 2007, 8:35–46.

9. Kleinjan DJ, van Heyningen V: Position effect in human genetic disease. Hum Mol Genet 1998, 7:1611–1618.

10. Sturtevant AH: The Effects of Unequal Crossing over at the Bar Locus in Drosophila. Genetics 1925, 10:117–147.

11. Chen X, Zhang J: The genomic landscape of position effects on protein expression level and noise in yeast. Cell Syst 2016, 2:347–354.

12. Chen M, Licon K, Otsuka R, Pillus L, Ideker T: Decoupling epigenetic and genetic effects through systematic analysis of gene position. Cell reports 2013, 3:128–137.

13. Dey SS, Foley JE, Limsirichai P, Schaffer DV, Arkin AP: Orthogonal control of expression mean and variance by epigenetic features at different genomic loci. Mol Syst Biol 2015, 11:806.

14. Batada NN, Hurst LD: Evolution of chromosome organization driven by selection for reduced gene expression noise. Nat Genet 2007, 39:945–949.

15. Wilson C, Bellen HJ, Gehring WJ: Position effects on eukaryotic gene expression. Annu Rev Cell Biol 1990, 6:679–714.

16. Milot E, Fraser P, Grosveld F: Position effects and genetic disease. Trends Genet 1996, 12:123–126.

17. Chen X, Zhang J: The Genomic Landscape of Position Effects on Protein Expression Level and Noise in Yeast. Cell Syst 2016, 2:347–354.

18. Pande A, Brosius J, Makalowska I, Makalowski W, Raabe CA: Transcriptional interference by small transcripts in proximal promoter regions. Nucleic Acids Res 2018, 46:1069–1088.

19. Strainic MG, Jr., Sullivan JJ, Collado-Vides J, deHaseth PL: Promoter interference in a bacteriophage lambda control region: effects of a range of interpromoter distances. J Bacteriol 2000, 182:216–220.

20. Pal C, Hurst LD: Evidence for co-evolution of gene order and recombination rate. Nat Genet 2003, 33:392–395.

21. Yang Y-F, Cao W, Wu S, Qian W: Genetic interaction network as an important determinant of gene order in genome evolution. Molecular biology and evolution 2017, 34:3254–3266.

22. Wang Z, Zhang J: Impact of gene expression noise on organismal fitness and the efficacy of natural selection. Proc Natl Acad Sci U S A 2011, 108:E67–76.

23. Artieri CG, Fraser HB: Evolution at two levels of gene expression in yeast. Genome Res 2014, 24:411–421.

24. Ghanbarian AT, Hurst LD: Neighboring Genes Show Correlated Evolution in Gene Expression. Mol Biol Evol 2015, 32:1748–1766.

25. Jackson DA, Hassan AB, Errington RJ, Cook PR: Visualization of focal sites of transcription within human nuclei. EMBO J 1993, 12:1059–1065.

26. Duan Z, Andronescu M, Schutz K, McIlwain S, Kim YJ, Lee C, Shendure J, Fields S, Blau CA, Noble WS: A three-dimensional model of the yeast genome. Nature 2010, 465:363–367.

27. Wang Y, Li X, Hu H: H3K4me2 reliably defines transcription factor binding regions in different cells. Genomics 2014, 103:222–228.

28. Kafri M, Metzl-Raz E, Jona G, Barkai N: The cost of protein production. Cell reports 2016, 14:22–31.

29. Dekel E, Alon U: Optimality and evolutionary tuning of the expression level of a protein. Nature 2005, 436:588–592.

30. Tan L, Xing D, Chang CH, Li H, Xie XS: Three-dimensional genome structures of single diploid human cells. Science 2018, 361:924–928.

31. Cherry JM, Hong EL, Amundsen C, Balakrishnan R, Binkley G, Chan ET, Christie KR, Costanzo MC, Dwight SS, Engel SR, et al: Saccharomyces Genome Database: the genomics resource of budding yeast. Nucleic Acids Res 2012, 40:D700–705.

32. Bolger AM, Lohse M, Usadel B: Trimmomatic: a flexible trimmer for Illumina sequence data. Bioinformatics 2014, 30:2114–2120.

33. Kim D, Paggi JM, Park C, Bennett C, Salzberg SL: Graph-based genome alignment and genotyping with HISAT2 and HISAT-genotype. Nat Biotechnol 2019, 37:907–915.

34. Wagner GP, Kin K, Lynch VJ: Measurement of mRNA abundance using RNA-seq data: RPKM measure is inconsistent among samples. Theory Biosci 2012, 131:281–285.

35. Pertea M, Pertea GM, Antonescu CM, Chang TC, Mendell JT, Salzberg SL: StringTie enables improved reconstruction of a transcriptome from RNA-seq reads. Nature Biotechnology 2015, 33:290.

36. Livak KJ, Schmittgen TD: Analysis of relative gene expression data using real-time quantitative PCR and the 2- ΔΔCT method. methods 2001, 25:402–408.

37. Murakami H, Ogata Y, Uchida S, Sasatomi T, Murakami N, Yamaguchi K, Gotanda Y, Akagi Y, Ishibashi N, Shirouzu K: [Therapeutic results of hepatic resection using thermal ablation for unresectable colorectal liver metastases]. Gan To Kagaku Ryoho 2009, 36:2039–2041.

